# l1kdeconv: an R package for peak calling analysis with LINCS L1000 data

**DOI:** 10.1101/165258

**Authors:** Zhao Li, Jin Li, Peng Yu

**Affiliations:** Department of Electrical and Computer Engineering, Texas A&M University, College Station, TX 77843, USA; TEES-AgriLife Center for Bioinformatics and Genomic Systems Engineering, Texas A&M University, College Station, TX 77843, USA

## Abstract

**Background:** LINCS L1000 is a high-throughput technology that allows gene expression measurement in a large number of assays. However, to fit the measurements of ~ 1000 genes in the ~ 500 color channels of LINCS L1000, every two landmark genes are designed to share a single channel. Thus, a deconvolution step is required to infer the expression values of each gene. Any errors in this step can be propagated adversely to the downstream analyses.

**Results:** We presented a LINCS L1000 data peak calling R package **l1kdeconv** based on a new outlier detection method and an aggregate Gaussian mixture model (AGMM). Upon the remove of outliers and the borrowing information among similar samples, **l1kdeconv** showed more stable and better performance than methods commonly used in LINCS L1000 data deconvolution.

**Conclusions:** Based on the benchmark using both simulated data and real data, the **l1kdeconv** package achieved more stable results than the commonly used LINCS L1000 data deconvolution methods.

## Introduction

The NIH Common Fund’ s Library of Integrated Network-based Cellular Signatures (LINCS) is a rich collection of gene expression data from a variety of human cell lines perturbed with a large battery of drugs and small molecules (Wang, Clark et al. 2016) (Lamb, Crawford et al. 2006). So far, data generated by LINCS L1000 technology comprise over 1,000,000 gene expression profiles from 42,553 perturbagens applied to as many as 77 cell lines. A total of 19,811 of the perturbagens (including over 2,000 FDA approved & clinical trial drugs) are small molecule compounds applied at different time points and doses. The other perturbagens are genetic perturbations, including knockdown and overexpression of well-selected 4,372 genes. In general, triplicated (or more) measurements were performed for each perturbergen, leading to a total of over 400,000 gene expression signatures generated by this technology (Subramanian, Narayan et al. 2017). Different from other genome wide expression measurement technologies such as microarray and RNA-Seq, L1000 is used in LINCS and only measures about 1000 selected “ landmark” genes with only about 500 distinct bead colors (Duan, Flynn et al. 2014) so that every two “ landmark” genes share a color. Despite this reduction greatly increasing the throughput of the perturbed assays that can be performed simultaneously and driving down the experimental cost, it induces an additional deconvolution analysis step that is crucial for the accuracy of the measure of the gene expression levels (Liu, Su et al. 2015).

This peak deconvolution step is necessary because each bead color is associated with two genes rather than one. For each color, deconvolution is supposed to yield a distribution that generally consists of two peaks. The default approach provided by LINCS is the *k*-medians clustering algorithm which partitions all the data into two distinct components, and the median expression values are assigned as the expression value of the two gene of a color. Liu *et al*. (Liu, Su et al. 2015) proposed another deconvolution approach that was based on naïve Gaussian mixture model (GMM) with only two components, and the expression values were inferred from the component means. However, although the two genes with the same color were selected so that their intensity differences are as much as possible, there still exist many cases in which the expressions of a pair of genes are intermixed, making it difficult to infer the expression values by either *k*-medians or naïve GMM (El-Melegy 2014). Thus, it is critical to develop a new method to improve the deconvolution accuracy.

Here, we presented an R package, **l1kdeconv**, that implements a new peak-calling algorithm based on an aggregate Gaussian mixture model (AGMM) along with an outlier detection method for improving the robustness of AGMM. AGMM is based on the Gaussian mixture model and borrows information from the samples of the same condition to improve the peaking calling accuracy.

## Implementation

### Outlier Detection in LINCS L1000 data

Gaussian mixture models based on clustering analysis methods in general are sensitive to outliers (Scott 2004). To improve the clustering accuracy, we first developed an outlier detection method to remove the outliers before peak calling.

Given the bead intensities of a color, we first estimated the kernel density of the bead intensities using the R function `density`(). When the estimated density is less than the threshold `dy_thr` (0.0001 as the default), the corresponding regions are recognized as data free gaps. Using these data free gaps, the beads are split into different clusters, as shown in **Figure 1**. Finally, if the size of a cluster is less than the threshold `clusterise_thr` (3 as the default), the beads in it will be identified as outliers. See **Figure 2** for the flowchart of the outlier detection.

**Figure 1.**
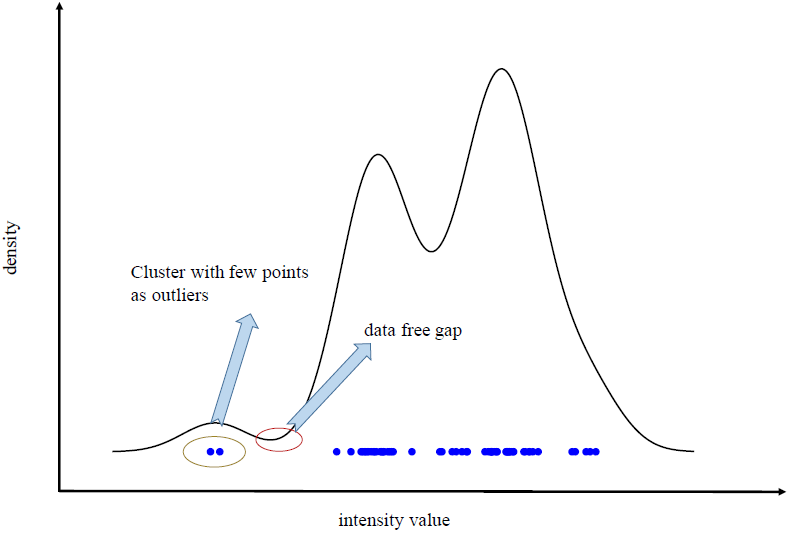
The overview of the outlier detection method in LINCS L1000 data. The curve represents the density function of the bead intensities of a color in a sample. The region in the red circle is a region in which there are no data points. This region splits the data into two regions. The region on the left contains too few data points, and the blue points circled in the brown eclipse are considered outliers as they degrade the performance of a 2-component Gaussian mixture model.

**Figure 2.**
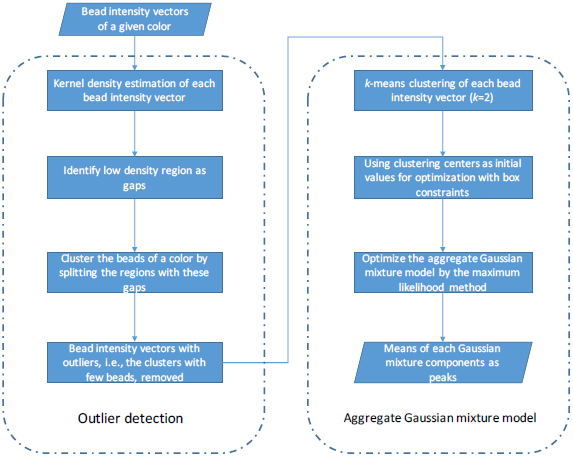
The implementation framework of **l1kdeconv** that includes the outlier detection and the aggregate Gaussian mixture model.

### aggregate Gaussian mixture model

To accurately detect the expression values of a pair of genes, we introduced a novel peak calling method to borrow the information of the samples at the same condition (e.g., the same cell line, the same perturbation and the same treatment duration). This method is an extension of naïve GMM with a constraint that the order of the gene expression values of two genes of the same color is consistent across these similar samples.

After the outliers were removed in the outlier-detection step, the two peaks of the pair of genes of a given color can be computed in the following way. We assumed the color intensities *X* of the beads measuring any given gene follow a Gaussian distribution 𝒩(*μ, σ^2^*,) with *μ* and *σ*^2^ as the mean and variance of bead intensities of the gene. Since each bead of a color can come from any of the two genes of the color, a two-component Gaussian mixture model was used to estimate the mean intensity of each gene. Due to the limited number of beads of each color, in order to have robust estimates of the intensity variance of each gene of a color, a common variance *σ*^2^ was assumed. Therefore, the probability density function of the bead intensities for a given color in a sample is

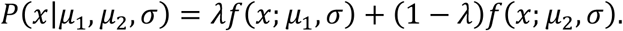

with the probability density function of a Gaussian distribution

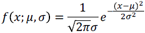

where *λ* is the proportion of the beads of the first gene and 1 - *λ* is the proportion of the beads of the second gene, *μ*_1_and*μ*_2_ are the mean intensities of the first and second genes respectively.

Because the two genes assigned to a color were selected in a way such that their intensity difference was maximized (Liu, Su et al. 2015), this difference between the two genes can be assumed to be in the same direction across the similar samples. In other words, of any given color, the mean intensity of one gene was always greater than the mean intensity of the other gene. This assumption suggested that the information borrowing among the similar samples will make the mean intensity estimates more robust.

The detailed method was described as below using an aggregate Gaussian mixture model (AGMM). Supposing there are m similar samples, we define the aggregate intensity vectors of a given color as {**x** _1_, **x**_2_, …, **x**_m_}, where each **x**_i_ = {*x*_i,1_, …, *x*_i,ni_} is a bead intensity vector of the color in the i-th sample and n_i_ is the number of beads. The log-likelihood of the AGMM is:

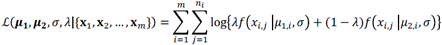

where *λ* is the proportion of the beads of the gene with a higher gene expression and 1 - *λ* is the proportion of the beads of the gene with a lower gene expression, ***μ***_1_ = {*μ* _1,1_, …, *μ*_1,m_} and ***μ***_2_ ={*μ* _2,1_, …, *μ*_2,m_} are the vectors of the two mean intensities in the *m* samples with the given color, and 03BB; is the standard deviation across these *m* samples with the given color. The maximum likelihood method can be used to estimate the parameters (***μ***_1_, ***μ***_2_, *σ* and *λ*) by optimizing the log-likelihood function of the AGMM. To enforce the differences between ***μ***_1_ and ***μ***_2_ of the same colors across similar samples having the same sign, we reparametrize the log-likelihood as follows so that box constraint optimization can be used:

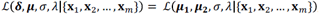

where δ and *μ* are defined as

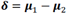

and

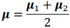

### Brief overview of the package

The **l1kdeconv** package contains a novel peak calling algorithm for LINCS L1000 data, using aggregate Gaussian mixture model, an extension of the naïve GMM. Rather than fitting the bead intensities of each color in a single sample using two-component Gaussian mixture model separately, AGMM calls the two peaks of each color across similar samples simultaneously. In addition, the outlier detection method included in **l1kdeconv** can also enhance the clustering performance of AGMM. **Figure 2** shows the framework of the outlier detection and AGMM approach.

The parameters of AGMM can be estimated using the R function optim()by the method of maximum likelihood. The L-BFGS-B method (Byrd, Lu et al. 1995) (Xiao and Wei 2007) is used to add box constrains. To ensure the convergence of the optimization, *k*-means is used to compute the rough estimates of the mean intensities, which are used as initial values for the optimization. To make the package user-friendly, detailed help pages and running examples are provided in the package.

## Results

### The construction of simulation dataset

To make a realistic evaluation of the performance of **l1kdeconv**, a simulated dataset was created with the key characteristics of LINCS L1000 data using a hierarchical model as described below. First, the empirical true expression values of each pair of genes were extracted by *k*-medians using the LINCS L1000 data of the A375 cell line treated with luciferase. Since the differences and averages of the empirical true expression values approximately followed a Gamma distribution and a Gaussian distribution respectively, these two distributions were used to resample the differences and the averages of the mean expression values of the two genes of any color. These differences and averages were then converted to the mean expression values of each pair of genes. The bead intensities were simulated from a two-component Gaussian mixture model with these mean expression values as the center of each Gaussian component. Then outliers were added to each color using a uniform distribution, and the number of outliers was generated from a Poisson distribution. Ten samples were created so that AGMM can borrow information across them. In each sample, there were 500 colors. In each color, there were roughly 60 beads and the ratio of the number of the beads for each gene in a color was 2:1.

### Comparison of l1kdeconv with *k*-medians and naïve GMM on the simulated dataset

Using the simulated data, a series of comparisons were conducted among *k*-medians, naïve GMM and AGMM with outlier detection in the **l1kdeconv** package. To have a fair comparison, we compared our AGMM with *k*-medians and naïve GMM that are also used in the standard LINCS L1000 deconvolution method. **Figure 3** shows the hexplot between true peaks and predicted peaks of each method. AGMM with outlier detection (D) is more accurate than *k*- medians (A) and naïve GMM (B). The Pearson Correlation Coefficients (PCC) are 0.76, 0.89 and 0.97 for *k*-medians, naïve GMM and AGMM with outlier detection, respectively. Then a test of significance for the difference between the two correlations based on dependent groups is conducted. The Williams’ s Test (Diedenhofen and Musch 2015) (Hittner, May et al. 2003) between the correlations shows that the PCC of AGMM with outlier detection is more significant than *k*-medians and naïve GMM with the *p*-value less than 1×10^-20^.

**Figure 3.**
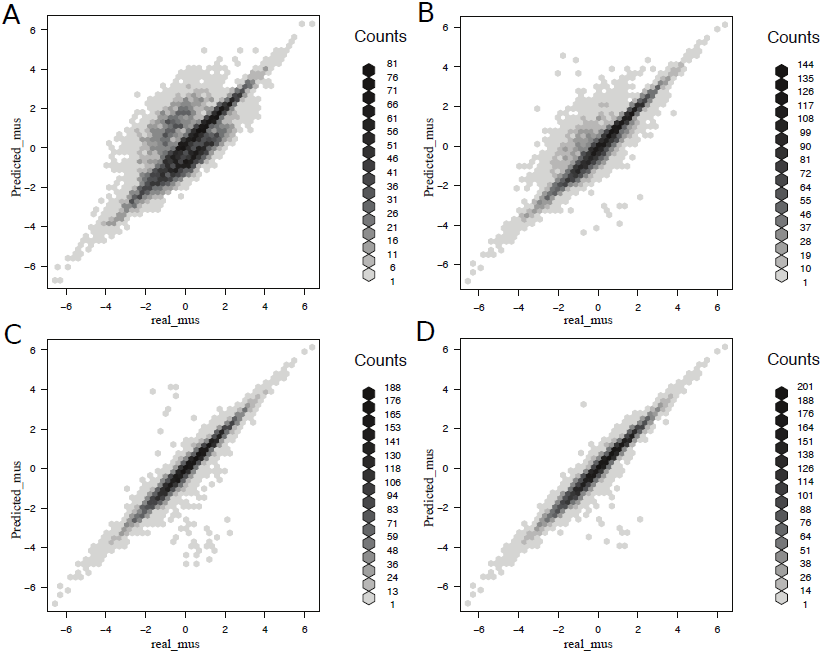
The comparison of the hexbin scatter plots of the intensity values of the true peaks and the predicted peaks among several methods. (A) *k*-medians (B) naï ve GMM, (C) AGMM without outlier detection and (D) AGMM with outlier detection. The darkness of the color of the hexagon indicates the number of points in it. The more points around y= x means the more accurate the prediction. AGMM with outlier detection outperforms *k*-medians, naï ve GMM and AGMM without outlier detection.

Comparisons were made among these methods with respected to the true prediction which is defined a prediction whose difference from the real value is less than 0.05. **Table 1** shows the number of the true predictions of each method and the corresponding mean absolute error (Hyndman and Koehler 2006). According to the numbers of true predictions, AGMM with the outlier detection resulted in 131% and 36% improvements compared with *k*-medians and naïve GMM, respectively. In addition, AGMM without outlier detection results in a reduction of 6%, indicating the importance of the outlier detection as a filtering step before calling AGMM.

**Table 1.**
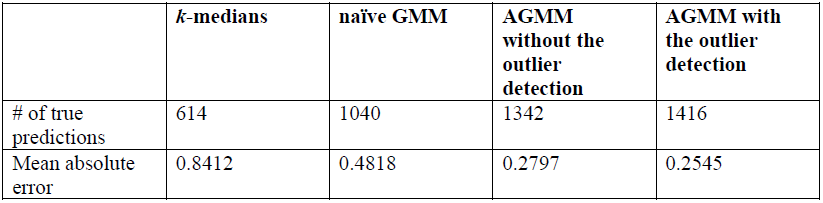
The number of the true predictions and mean absolute error of *k*-medians, naïve GMM, AGMM without the outlier detection and AGMM with the outlier detection. AGMM with the outlier detection made more true predictions, and its predictions have the least mean absolute error.

|  | **$k$-medians** | **naïve GMM** | **AGMM**<br>**without the**<br>**outlier**<br>**detection** | **AGMM with**<br>**the outlier**<br>**detection** |
| --- | --- | --- | --- | --- |
| # of true predictions | 614 | 1040 | 1342 | 1416 |
| Mean absolute error | 0.8412 | 0.4818 | 0.2797 | 0.2545 |

To further benchmark the accuracy of the three methods, we compared whether the order of the predicted two peaks were flipped. **Figure 4** shows that AGMM with outlier detection indeed greatly improves the performance with respect to the flipping errors because of its ability to borrow information across similar samples. In contrast, *k*-medians and naïve GMM lack this ability, thus they produced many flipping cases.

**Figure 4.**
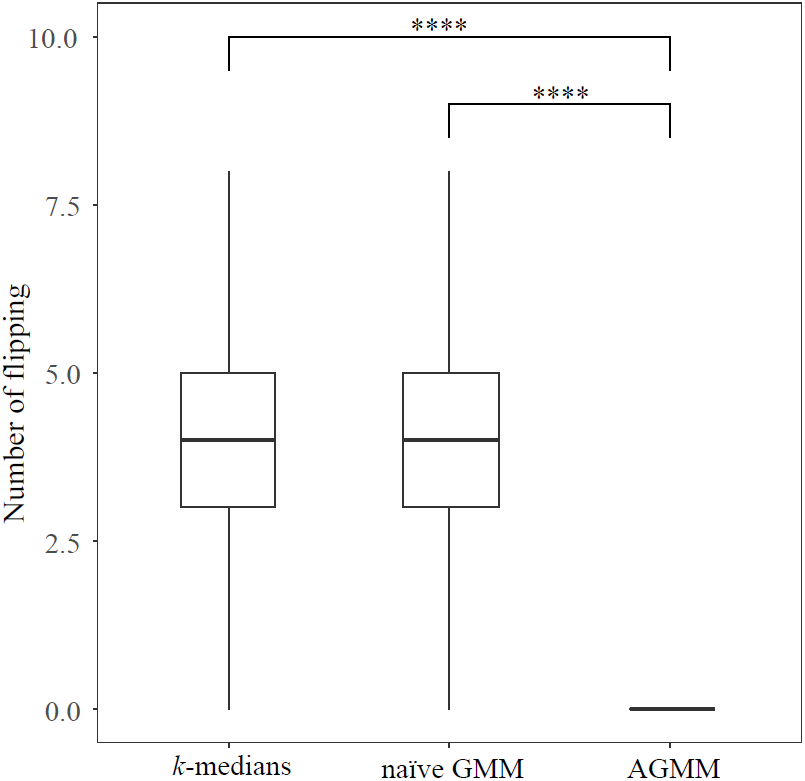
The boxplot of the numbers of flippings in each color across similar samples of *k*- medians, naïve GMM and AGMM with outlier detection. AGMM with outlier detection produces no flipping in this simulated dataset. The last method significantly *p* outperforms the first two methods (**** means< 0.0001).

### Comparison of l1kdeconv with *k*-medians and naïve GMM on a real dataset

The comparison using the simulated data confirmed that AGMM with outlier detection is more stable and more accurate than *k*-medians and naïve GMM. To demonstrate the performance improvement of **l1kdeconv** on real data, we used a dataset of A375 cells treated with luciferase from LINCS L1000 Phase II data (GSE70138). Eleven replicates were used to borrow information across these replicates. Unlike calling the peaks of each replicate separately by *k*- medians and naïve GMM, AGMM filters the outlier of each sample, and then deconvolutes them in one group. **Figure 5** shows nine representative samples from a color in this real dataset. *k*- medians (**a** and **c**) and naïve GMM (**b**) were unable to identify the two peaks correctly as the order of the two peaks are flipped. However, AGMM with outlier detection can identify the two peaks correctly because the information in the easy cases of **d** - **i** can be used to help with the deconvolution of the difficult ones (**a**, **b** and **c**). In the cases **d**, **e** and **f**, although the results from *k*-medians and naïve GMM are not flipped, AGMM with the outlier detection can fit the real data better. The peak coordinates predicted by *k*-medians, naïve GMM and AGMM with outlier detection in **Figure 5** are shown in **Table 2**.

**Table 2.** The deconvolution results of *k*-medians, naï ve GMM and AGMM with outlier detection using real data. Each pair of superscripts indicates a peak flipping case. The row index refers to the subfigure label in Figure 5.

| | $k$-medians | | naïve GMM | | AGMM with outlier detection | |
| --- | --- | --- | --- | --- | --- | --- |
|  | Peak1 | Peak2 | Peak1 | Peak2 | Peak1 | Peak2 |
| a | -0.32<sup>1</sup> | -0.09<sup>1</sup> | -0.58 | -0.21 | -0.43 | -0.19 |
| b | -0.29 | -0.11 | -0.21<sup>2</sup> | -0.10<sup>2</sup> | -0.27 | -0.16 |
| c | -0.06<sup>3</sup> | -0.37<sup>3</sup> | -0.43 | -0.17 | -0.41 | -0.20 |
| d | -0.60 | -0.21 | -1.95 | -0.36 | -0.65 | -0.23 |
| e | -0.69 | -0.20 | -0.70 | -0.23 | -0.64 | -0.22 |
| f | -0.69 | -0.11 | -0.55 | -0.07 | -0.63 | -0.09 |
| g | -0.72 | -0.22 | -0.78 | -0.22 | -0.75 | -0.20 |
| h | -0.78 | -0.16 | -0.79 | -0.14 | -0.80 | -0.14 |
| i | -0.70 | -0.15 | -0.77 | -0.17 | -0.76 | -0.15 |

**Figure 5.**
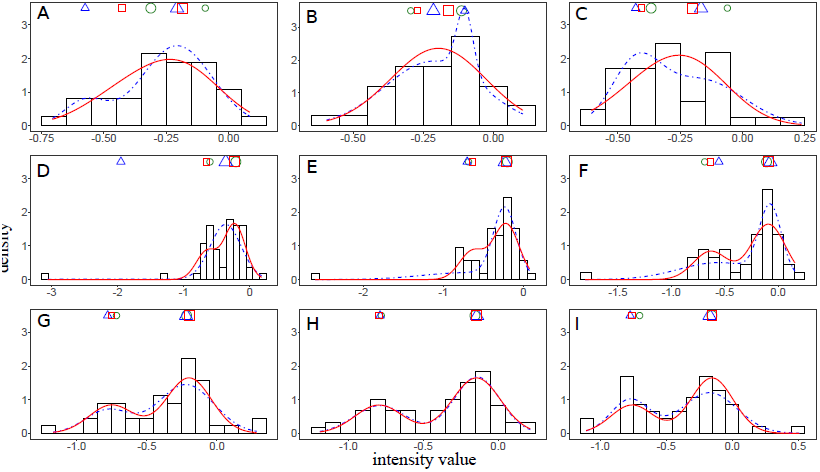
The performance comparison of *k*-medians, naï ve GMM and AGMM with outlier detection. Subfigures (a - i) show the intensity histograms of a color of A375 cell line samples treated with luciferase. The blue curves and red curves represent the density functions estimated by naï ve GMM and AGMM, respectively. The two cluster centers are indicated by the green, blue and red shapes above the histogram learned by *k*-medians, naï ve GMM and AGMM with outlier detection, respectively. The bigger/smaller shapes indicate the cluster with more/less beads.

## Conclusions

The **l1kdeconv** package provides a stable and accurate deconvolution algorithm for LINCS L1000 data. Because the deconvolution is the first step of the analysis of LINCS L1000 data, any significant improvement in this step will have a critical impact on the downstream analyses. The **l1kdeconv** package has two components --- the outlier detection and aggregated Gaussian mixture model. By filtering the outliers and borrowing information from similar samples, **l1kdeconv** outperforms *k*-medians and naïve Gaussian mixture model using the limited number of intensity values.

The package also provides detailed help pages, which makes it user friendly. Furthermore, we standardize the distribution, installation and maintenance of this package. It is available on the Comprehensive R Archive Network (CRAN) at http://cran.r-project.org.

## Declarations

### Ethics approval and consent to participate

Not applicable.

### Consent to publish

Not applicable.

### Availability of data and materials

The **l1kdeconv** package is available on the Comprehensive R Archive Network (CRAN) at http://cran.r-project.org. The LINCS L1000 data is available at GEO with the accession number of GSE70138.

### Authors’ Contributions

PY conceived the general project and supervised it. ZL and PY were the principal developers. ZL, JL and PY wrote the manuscript. All the authors contributed with ideas, tested the software, read the final manuscript and approved it.

## Acknowledgements

The study was supported by the startup funding to Peng Yu from the ECE department and Texas A&M Engineering Experiment Station/Dwight Look College of Engineering at Texas A&M University and by funding from TEES-AgriLife Center for Bioinformatics and Genomic Systems Engineering (CBGSE) at Texas A&M University.

